# Infection Groups Differential (IGD) Score Reveals Infection Ability Difference between SARS-CoV-2 and Other Coronaviruses

**DOI:** 10.1101/2020.05.12.090324

**Authors:** Ziwei Song, Xingchen Zhou, Yuanyuan Cai, Shuo Feng, Tingting Zhang, Yun Wang, Maode Lai, Jing Li

**Affiliations:** State Key Laboratory of Natural Medicines, China Pharmaceutical University, Nanjing 210009, China; School of Life Science and Technology, China Pharmaceutical University, Nanjing 210009, China; School of Traditional Chinese Pharmacy, China Pharmaceutical University, Nanjing 210009, China

## Abstract

The Corona Virus Disease 2019 (COVID-19) pandemic that began in late December 2019 has resulted in millions of cases diagnosed worldwide. Reports have shown that SARS-CoV-2 shows extremely higher infection rates than other coronaviruses. This study conducted a phylogenetics analysis of 91 representative coronaviruses and found that the functional spike protein of SARS-CoV-2, which interacts with the human receptor ACE2, is actually not undergoing distinct selection pressure compared to other coronaviruses. Furthermore, we define a new measurement, infection group differential (IGD) score, in assessing the infection ability of two human coronavirus groups. There are nine extremely high IGD (ehIGD) sites in the receptor-binding domain (RBD) out of 40 high IGD (hIGD) sites that exhibit a unique infection-related pattern from the haplotype network and docking energy comparison. These 40 hIGD sites are basically conserved among the SARS-CoV-2, i.e. there are only two hIGD sites mutated in four out of 1,058 samples, defined as rare-mutation hIGD (rhIGD) sites. In conclusion, ehIGD and rhIGD sites might be of great significance to the development of vaccines.

## Introduction

The COVID-19 pandemic caused by the new coronavirus (SARS-CoV-2) continues to spread around the world as of this writing [1]. It is having a broad and profound impact on the global health, politics, economy, and society [2]. Apparently, SARS-CoV-2 has caused more infections and deaths than SARS coronavirus (SARS-CoV) and MERS coronavirus (MERS-CoV), whereas current research indicates that these all originated from bats [3, 4]. Therefore, it is extremely important to investigate the mutation and evolutionary characteristics of SARS-CoV-2, and the analysis of the structure that causes its strong infectivity is of great significance to the development of vaccines and control of the pandemic.

Coronaviruses are RNA viruses that consist of four subtypes: α, β, γ, and δ [5]. SARS-CoV-2 belongs to the family of β coronaviruses and is the seventh known coronavirus that can infect humans [6]. The remaining six human coronaviruses are HCoV-229E, HCoV-OC43, HCoV-NL63, HCoV-HKU1, SARS-CoV (causes Severe Acute Respiratory Syndrome) and MERS-CoV (triggers Middle East Respiratory Syndrome) [7]. Although the above coronaviruses can all infect humans, studies have revealed different pathogenicity and variable clinical manifestations. Some coronaviruses (HCoV-229E, HCoV-OC43, HCoV-NL63, and HCoV-HKU1) only cause common cold, whereas SARS-CoV (R0 = 2-5, case fatality rate = 10%), MERS-CoV (R0 = 0.3-0.8, case fatality rate = 40%), and SARS-CoV-2 can cause fever, cough, and pneumonia, which can be fatal in severe cases [8].

The spike protein on the SARS-CoV-2 surface plays a key role in the invasion into human cells [9]. Its structure determines whether it can infect humans and how capable it is. First, the restriction sites on the spike protein determine how the coronavirus is packaged into human cells. The spike protein binds to the angiotensin-converting enzyme 2 (ACE2) on human cells through the receptor-binding domain (RBD), which then initiates the infection [10]. Therefore, these restriction sites and RBD characteristics have always been the focus of coronavirus research studies. Therefore, the spike protein is of vital importance in studying the infectivity and virulence of SARS-CoV-2 [11, 12].

Recent research indicates that there is a polybasic cleavage site at the junction of the S1 and S2 subunits of the spike protein of SARS-CoV-2, namely the Furin protease cleavage site (RRAR), and formed a special O-linked glycan structure [9]. In addition, a comparison study of SARS-CoV and SARS-CoV-2 has shown that six amino acid residues in RBD are the key sites for binding to human ACE2. However, five of them in the SARS-CoV-2 sequence have been mutated, which may be responsible for the enhanced binding to human ACE2 [13]. The abovementioned restriction sites and mutations in the RBD structure have always been the research hotspots of SARS-CoV-2. Currently, most of the functional key residues were identified by direct interactions, i.e., by exploring the protein-protein docking sites or enzyme-substrate binding sites. In this work, we identify the sites that may be associated with infection ability through sequence comparisons, as well as determine significant variants between human and non-human coronaviruses. The functional effects of these sites were evaluated by the protein-protein virtual docking energy, and we plan to further assess these in a cell model in our following study.

## Methods

### Data acquisition

We downloaded all of the SARS-like coronaviruses that were more than 5,000 bp in length from GenBank via VIPR (https://www.viprbrc.org/brc/vipr_genome_search.spg?method=ShowCleanSearch&decorator=corona, 2020/01/25). The metadata and URLs of these sequences are shown in Table S1. The reference sequence of SARS-CoV-2 was downloaded from GenBank (https://www.ncbi.nlm.nih.gov/nuccore/MN908947.3?report=genbank, 2020/01/25). Approximately 44 spike proteins predicted from these coronavirus genome data were used in searching for homologous sequences in the database. Homologous sequence searching of SARS-like coronaviruses spike protein was first performed by BLASTP (v 2.2.29+) [14] in NR (Non-Redundant Protein Sequence Database), and 868 sequences were selected under the condition of sequence length > 600 aa and a sequence identity > 30%. Next, the 868 sequences were classified into 124 clusters criteria of sequence identity ≥ 95% and sequence length ≥ 90%. Taking the longest sequences as the representative strain in each cluster and checking the NCBI genome database, 91 of 124 strains were found to have complete genome sequences in NCBI. Table S2 lists the accession numbers and taxonomic characteristics of 91 representative coronaviruses. Finally, the protein sequences of each coronavirus were predicted by GeneMarks from the genome DNA sequences and used in the subsequent phylogenetic analysis.

1458 SARS-CoV-2 genomes were downloaded from the GISAID database (https://www.gisaid.org/, 2020/03/25). After removing the non-human strains and strains with low sequence quality, 1,058 SARS-CoV-2 strains were used in the phylogenetic analysis.

### Phylogenetic analysis

The DNA sequences of whole genome and related protein sequences of 91 coronaviruses were aligned using mafft v7.455 [15], and the result multiple sequence was trimmed for poorly aligned positions with Gblock 0.91b [16]. RAxML v8.2.12 [17] was used to build the maximum likelihood phylogenetic tree of genomes with the parameters “-m GTRCAT” and protein with the parameters “-m PROTGAMMAILGX”. R package “ggtree” [18] was used to display phylogenetic tree.

### Codon usage bias analysis

Condonw v1.3 [19] was used to calculate the universal index value of each codon of the Cds of coronavirus functional protein.

### Ka/Ks analysis

Ka/KS ratios were calculated using KaKs_Calculator 2.0 [20] and used in the analysis of selection pressure.

### Visual analysis of multiple sequence alignments

Mafft was used to generate the multi-sequence aligned data-based amino acid sequence of coronavirus spike proteins and domains. R package “ggmsa” was used to visualize the results. From the multiple alignment, the sequence identities within and between groups could be calculated at each position. The “Ratio (3 & 83)” were calculated from the identities between the high infection human coronaviruses group and none-human coronaviruses group. The “Ratio (5 & 83)” were calculated from the identities between the low infection human coronaviruses group and none-human coronaviruses group. The infection group (similarity) differential (IGD) scores were calculated from ratio (3 & 83) and ratio (5 & 83). From the total 1,273 positions of S1 subunit of spike protein, the mean and sd values were calculated, thus the high IGD (hIGD) sites defined as IGD score greater than the mean + 3sd value, while the extremely high IGD (ehIGD) site defined as IGD score greater than the mean + 5sd value. For each ehIGD in S1 subunit, the hIGD and ehIGD sites in the range of up- and down-stream 25 amino acids residues were connected and defined as a hIGD region. Furthermore, the rhIGD sites defined as the rare mutation with frequency less than 0.5% of hIGD sites among SARS-CoV-2 sequences.

### Haplotype network analysis

DnaSP v6.12.03 [21] was used to generate multi-sequence aligned haplotype data. Arlequin v3.5.2.2 [22] was used to estimate haplotype frequency. PopART v1.7 [23] was used to generate haplotype networks based on the haplotypes generated by DnaSP and Arlequin.

### Structure prediction and protein-protein docking

The structure of human ACE2 receptor was obtained from PDB database (https://www.rcsb.org/, PDB_ID: 6acc) [24]. The structure of the RBD domain of SARS-CoV-2 spike protein was obtained from PDB database (PDB_ID: 6m17) [25]. The structures of mutants of RBD for the ehIGD sites changing to the amino acids of other highly infectious coronavirus were homology modeling using SWISS-MODEL [26].

The structure of SARS-CoV-2 spike protein was homology modeling from the structure of SARS-CoV spike protein (PDB_ID: 6vsb) since it hasn’t been fully resolved yet till the submitted data of this work. Thus, the structures of mutants of spike protein for the rhIGD sites changing to the amino acids of other type of SARS-CoV-2 were also homology modeling using SWISS-MODEL.

For protein-protein docking, we employed the online software SwarmDock [27] and MOE (v2019) to perform the virtual docking between human ACE2 receptor and RBD domain, spike protein and mutants of SARS-CoV-2. For each complex of docking, we conducted 100 independent conformations, and selected the conformation with the lowest binding energy as the final result.

## Results

### Phylogenetic analysis of coronaviruses and SARS-CoV-2

The overview of this work is presented in Figure 1A. We searched all coronaviruses with spike protein in the NCBI database, which classified these into clusters and selected representative strains with complete genome sequences to avoid data collection bias.

**Figure 1.**
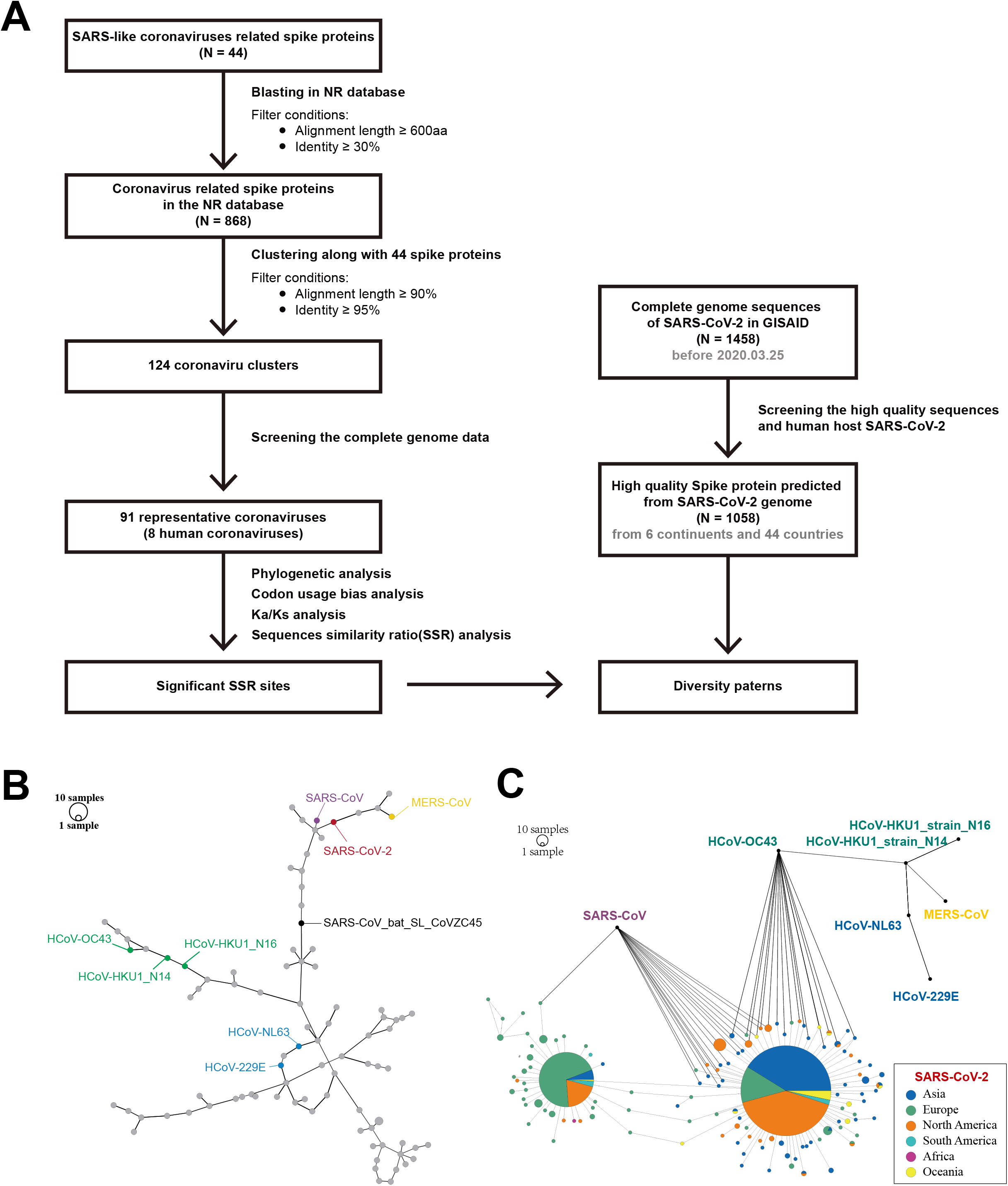
Phylogenetic comparisons of SARS-CoV-2 and other coronaviruses. (A) The workflow of this work; (B) The haplotype networks of RBD of 91 coronavirus genomes, and the representative sequence of SARS-CoV-2 here is “Wuhan-hu-1”, the detailed information is shown in Table S3; (C) The haplotype networks of spike proteins of 1,058 representative SARS-CoV-2 from around the world cases. Human coronaviruses are indicated by different colors, coronaviruses with middle infection ability are highlighted in green, and ones with lower infection ability are shown in blue. RBD, receptor binding domain.

Phylogenetic analysis of the entire genome of 91 representative coronaviruses shows that SARS-CoV-2 and SARS-CoV_bat_SL_CovZC45 have the closest evolutionary distance compared to other human coronaviruses (Figure S1). This result has been reported before, indicating that SARS-CoV-2 may have evolved from bat coronaviruses [28]. From Figure 1B, the haplotype network constructed based on the receptor-binding domain (RBD) of 91 representative coronavirus spike proteins, which showed that the haplotype of SARS-CoV-2 is on the same branch as the haplotype of SARS-CoV and MERS-CoV. On the other hand, the low infection human coronavirus, i.e. HCoV-229E, HCoV-OC43, HCoV-NL63, and HCoV-HKU1, cluster closely in the other branch. Intriguingly, the SARS-CoV_bat_SL_CoVZC45 located in the medium of high and low infection coronaviruses, unlike the phylogenetic relationship of whole genome presented in Figure.S1, indicating that the haplotype of the coronavirus RBD has a higher correlation to pathogenicity than phylogenetic relationship.

Simultaneously, we collected all available SARS-CoV-2 sequencing data from the GISAID database. After filtering nonhuman strains and low-quality sequencing strains, approximately 1,058 strains from total 43 countries of 6 continents remained (Table S4 and S5). Multiple sequence alignment of the S1 subunit of the global SARS-CoV-2 spike protein shows that the N-terminal domain (NTD) and RBD are conserved (identity > 97% within global SARS-CoV-2, Figure S2), particularly SARS-CoV-2 from the Asia RBD area (identity > 99%, Table S6). As shown in Figure 1C, the haplotype network analysis of these representative SARS-CoV-2 strains and 7 human coronaviruses shows that the spike protein haplotype of SARS-Cov-2 were clearly divided into two clusters, and closest to the haplotype of SARS-CoV. The larger cluster, which is closer to both SARS-CoV and HCoV-OC43, contain 768 haplotypes of SARS-CoV-2 composed by 42% Asian, 39% North American, and 14% European samples. And another cluster contains 290 haplotypes including 74% European, 17% North American, and 5% Asian samples. From this network, the haplotype of HCoV-OC43 is closer to SARS-Cov-2 haplotype than MERS-CoV. Thus, the haplotype network constructed from spike protein might reflect partial pathogenicity relationship but the specific sites with more influence on the infection need to be further explored.

### Selection analysis of coronaviruses and SARS-CoV-2

The main functional proteins of coronaviruses include the membrane protein, nucleocapsid protein, orf1a polyprotein, orf1b polyprotein, and spike protein. To explore the evolutionary characteristics of SARS-CoV-2 compared with other coronaviruses, we first compared the sequence identities of the entire genome and 5 functional proteins of 91 coronaviruses. The results show that the sequence of the orf1b polyprotein is relatively conserved, whereas the spike protein sequence differs among these coronaviruses (Figure S3). Then, we conducted codon preference analysis of different protein domains of the 91 coronaviruses, including spike protein, connection domain (CD), central helix (CH), heptad repeat 1 (HR1), NTD, and RBD. The results show that the codon preference of SARS-CoV-2 is basically similar to the other coronaviruses (Figure S4). Furthermore, Ka/Ks analysis of 5 functional proteins of 91 coronaviruses shows that the 5 functional proteins of SARS-CoV-2 mainly underwent neutral evolution in 91 coronaviruses (Figure S5A). However, the orf1a polyprotein gene of SARS-CoV-2 has been subjected to purification selection in human coronavirus, whereas the others underwent neutral evolution (Figure S5B). These findings suggest that the spike protein of SARS-CoV-2 did not undergo special selection pressure compared to other coronaviruses.

### Infection group differential (IGD) sites and regions in spike protein of SARS-CoV-2

To study sites of the spike protein of SARS-CoV-2 that are related to infectivity, we firstly classified 91 representative coronaviruses into three groups, i.e. high infection coronaviruses group (including SARS-Cov-2, SARS-CoV, and MERS-CoV), low infection coronaviruses group (including HCoV-229E, HCoV-OC43, HCoV-NL63, and HCoV-HKU1), none-human coronaviruses group (including 83 non-human coronaviruses). After that, we performed sequence alignment on 91 representative coronaviruses and calculated the similarity of each amino acid position within groups and between groups, as shown in Figure 2A, Figure 2B, Figure 2D and Figure 2E. Thus, the “Ratio (3 & 83)” and “Ratio (5 & 83)” should reflect the sequences differentiation between the human and none-human coronaviruses, as shown in Figure 2C and Figure 2F.

**Figure 2.**
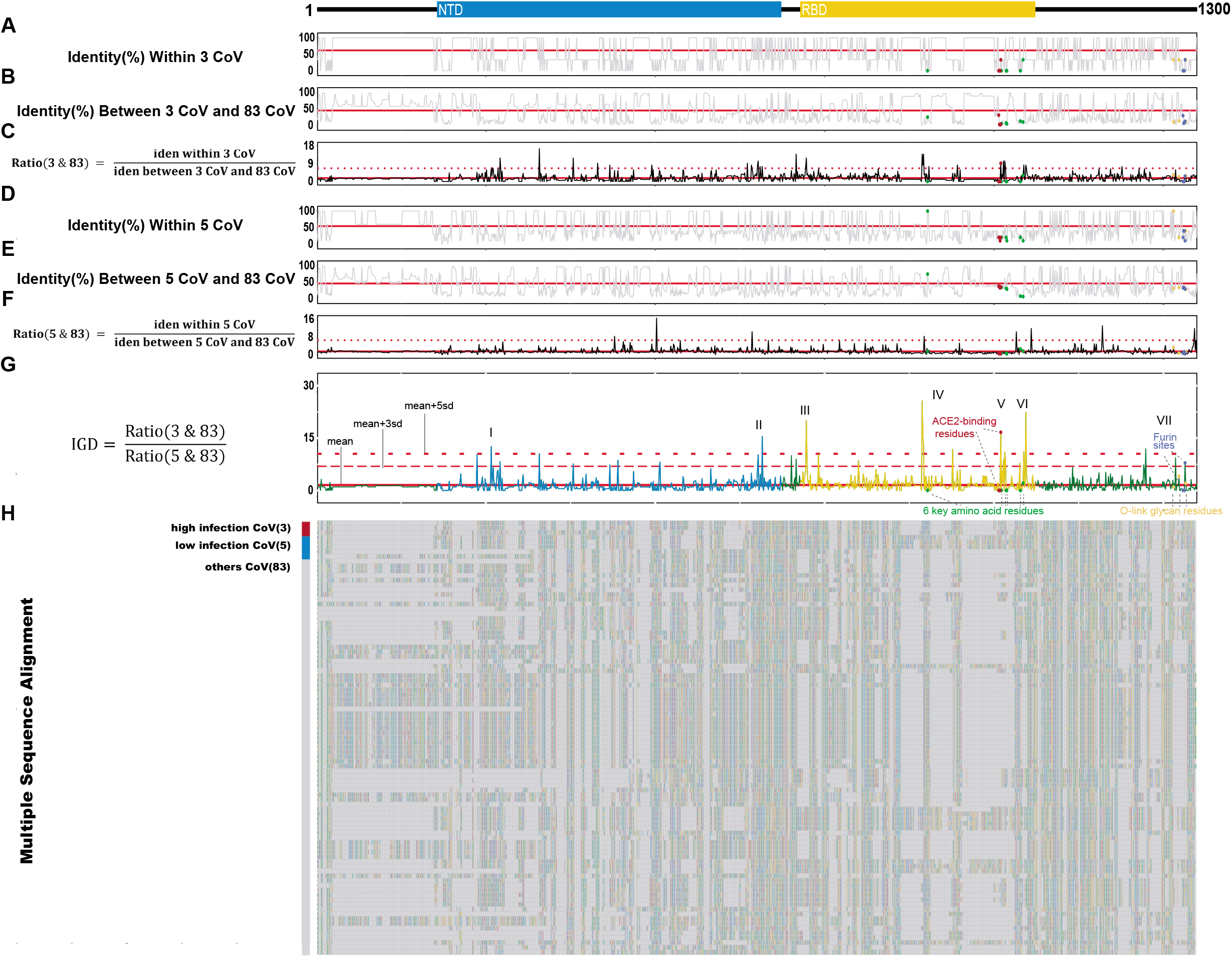
Multiple sequence alignment and identity pattern visualization of 91 representative coronaviruses spike protein S1 subunit. (A)-(C) Three curve graph compare the amino acid difference between human coronaviruses with high infection ability (SARA-CoV-2, SARS-CoV and MERS-CoV) and non-human coronaviruses; (D)-(F) Three curve graph compare the amino acid difference between human coronaviruses with lower infection ability (HCoV-OC43, HCoV-HKU1 (N = 2), HCoV-229E, and HCoV-NL63) and non-human coronaviruses. (G) The IGD score curve graph, indicate the amino acid residue differences between two groups of human coronaviruses with different infection ability; (H) The multiple sequence alignment of S1 subunit of 91 representative coronaviruses. The solid horizontal line represents the mean value of IGD score, the dashed horizontal line indicates the mean + 3sd value of IGD score, the dotted horizontal line depicts the mean + 5sd value of IGD score. Roman numerals indicate the region of the IGD sites. The specific sites previously reported is highlighted, i.e. the red dots indicate ACE2-binding residues, green dots indicate six key amino acid residues, purple dots indicate Furin cleavage sites, and yellow dots indicate O-linked glycan residues. IGD, infection group differential; CoV, coronavirus; NTD, N-terminal domain; RBD, receptor-binding domain.

Finally, we defined infection group differential (IGD) score to measure the residues feature between the high infection and low infection coronaviruses groups. As shown in Figure 2G, the IGD score of each position is calculated from the similarity ratio between the two groups of human coronaviruses. The high IGD (hIGD) score demonstrates the significant differentiation (IGD score greater than mean + 3sd value) between two human coronaviruses groups. Therefore, 40 hIGD sites in the spike protein sequence indicate the characteristic regions of coronavirus infectivity, shown in Figure S6 and Table S7a. In addition, we have defined the “Region” of hIGD sites in S1 subunit, thus the seven regions distributed with 31 hIGD sites. In particular, the six key residues of RBD binding to ACE2 [13] were assigned to Region IV, V and VI, while the Furin protease cleavage sites and the special O-link glycan residues [29] were located in Region VII, as shown in Figure 2G and Table S7a.

### Special hIGD sites and influence to SARS-Cov-2 infection

To further explore the potential importance of hIGD sites, we focus on some special hIGD sites, i.e. extremely high IGD (ehIGD) sites in RBD region and rhIGD sites from current dataset. Here, the ehIGD sites defined as IGD score greater than mean + 5sd value, and rhIGD indicated mutation of 40 hIGD in 1058 SARS-Cov-2 which are actually rare mutation with frequency less than 0.5%.

The specific positions of nine ehIGD sites and two rhIGD sites in S1 subunit of SARS-CoV-2 are shown in Figure 3A. The haplotype network of nine ehIGD sites in RBD is shown in Figure 3B. It shows that the distances between the haplotype of SARS-CoV-2 and other human coronaviruses are closely related to their infection ability. In particular, the most closely related coronaviruses are SARS-CoV and MERS-CoV, and there is only one differential amino acid between SARS-CoV-2 and SARS-CoV (SARS-CoV-2: G485; SARS-CoV: A485). However, the nine corresponding positions of HCoV-OC43, HCoV-HKU1_N14, and HCoV-HKU1_N16 are different from SARS-CoV-2. HCoV-229E and HCoV-NL63 did not even exist in the haplotype network because these have a total of nine deletion sites from the multiple alignments.

**Figure 3.**
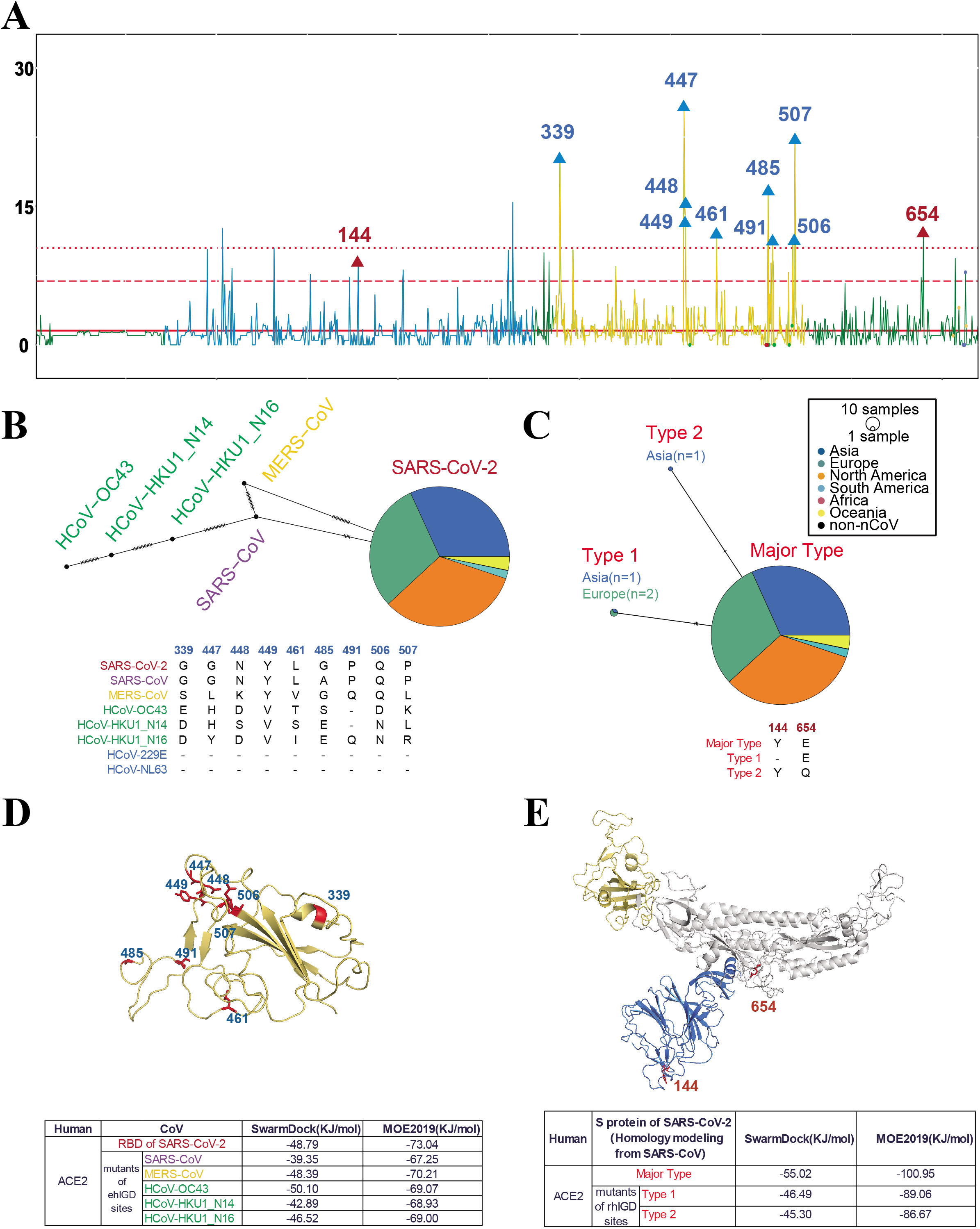
Haplotype analysis and molecular docking of ehIGD sites and rhIGD sites. (A) The IGD curve of S1 subunit of coronaviruses with different infection ability. The solid horizontal line represents the mean value of IGD score, the dashed horizontal line indicates the mean + 3sd value of IGD score, the dotted horizontal line depicts the mean + 5sd value of IGD score. hIGD, the IGD score is greater than the mean + 3sd value; ehIGD, the IGD score is greater than the mean + 5sd value; rhIGD sites, rare mutation hIGD sites among SARS-CoV-2 sequences from human. The ehIGD sites and rhIGD sites are marked by blue triangles and red triangles, respectively. The red dots indicate ACE2-binding residues, green dots indicate six key amino acid residues, purple dots indicate Furin cleavage sites, and yellow dots indicate O-linked glycan residues; (B) The haplotype network of nine ehIGD sites in RBD of 1,058 SARS-CoV-2 strains and other human coronaviruses; detailed information is shown in Table S9a; (C) The haplotype network of two rhIGD sites of 1,058 SARS-CoV-2 strains; The detailed information of ehIGD sites and rhIGD sites are shown in Table S9b; (D) Protein structure of SARS-CoV-2 RBD. The blue numbers and red sticks indicate the ehIGD sites. The following table shows the binding energies between different mutations of coronavirus and the human ACE2 receptor. (E) Protein structure of SARS-CoV-2 spike protein. The red numbers and red sticks indicate two rhIGD sites of S1 subunit. The following table shows the binding energies among the three types of SARS-CoV-2 and the human ACE2 receptor.

Among the total 40 hIGD sites from our analysis, only two rhIGD sites among the 1,058 SARS-CoV-2 strains actually occurred in only two Asian (India and Georgia) and two European (Netherlands) samples (Figure 3C, Table. S8). Compared with the majority of haplotype, the Type 1 haplotype has a deletion of residue 144, whereas the Type 2 haplotype has a substitution of E654 to Q654.

To verify the above crucial sites of SARS-CoV-2, we compared the affinity of the spike protein and ACE2 before and after ehIGD and rhIGD site mutation based on protein-protein docking (Figure 3D and 3E). The results show that the binding energy of SARS-CoV-2 and ACE2 is the lowest before mutation, especially when using MOE2019 for docking. Then the binding energy is basically increased after ehIGD sites are mutated (Figure 3D). Moreover, mutations at two rhIGD sites, which increases the binding energy, also seem to have an impact on the ability of SARS-CoV-2 to bind to ACE2 (Figure 3E). Therefore, nine ehIGD and two rhIGD sites of the spike protein are highly likely to be closely related to the high infectivity of SARS-CoV-2. However, additional investigations are warranted.

## Discussion

Coronaviruses mainly originate from animals, and some gradually evolve to infect humans. The different sequence characteristics of these human coronaviruses determine the differences in infection ability and pathogenicity. The high infection ability of SARS-CoV-2 is responsible for the rapid spread of COVID-19, and thus we attempted to study the distinctive sequence features of SARS-CoV-2.

We started with the spike protein of SARS-like coronaviruses and obtained 91 representative coronaviruses (including eight coronaviruses that can infect humans) through genome sequence alignment and screening. We analyzed the evolutionary characteristics of these coronaviruses and their spike proteins, and found that spike protein variations associated with receptor binding are the largest and mainly underwent neutral evolution among the functional proteins of these coronaviruses.

In this work, we defined a new measurement, namely, the sequence similarity ratio, to measure the sequence feature among coronavirus groups with different infection abilities. The haplotype networks constructed by the ehIGD residues reveal that the network distance between different coronaviruses strains and SARS-CoV-2 is proportional to their infection ability but not their phylogenetic distance. Therefore, we hypothesized that ehIGD residues influence the infection ability of SARS-CoV-2.

Only two rhIGD sites over 40 hIGD sites are not statistically significant (Fisher test, *p*=0.7653) compared with the total of 1058 polymorphic sites over 1,273 residues in the spike protein (Table S7b). These findings suggest that SARS-CoV-2 has not undergone large-scale major mutations yet, and the development of related vaccines are still important and of great significance.

Some of these crucial residues are also consistent with the functional sites in previous reports. For example, G482, V483, E484, and G485 are reported to be the functionally important epitopes of SARS-CoV-2 binding to ACE2 [11]. Another example is that SARS-CoV-2 has a unique ACE2 interacting residue K417 that forms salt-bridge interactions with ACE2 D30 [12]. G485 is in the list of our ehIGD sites, and we additionally found novel G339, G447, N448, Y449, L461, P491, Q506 and P507 residues in RBD with high IGS score between different infection ability coronaviruses. Whether these sites are related to the SARS-CoV-2 infection ability is unclear, and we are currently verifying this using a cell model.

The discovery and confirmation of key sites related to SARS-CoV-2 infection ability are crucial to the design of vaccines and therapeutic drugs. At current stage, the effect of ehIGD and rhIGD sites have been verified by virtual protein-protein docking. The future biologic evaluation is performing at cell level.

## Authors’ contributions

JL and ML conceived, designed the study and revise the manuscript. XZ, YC, SF, TZ and YW collected data and performed the analyses. JL and ZS interpreted the data and wrote the draft manuscript.

