## Supplementary figures and images for "Infection Groups Differential (IGD) Score Reveals Infection Ability Difference between SARS-CoV-2 and Other Coronaviruses"

### Supplementary Figures.pdf

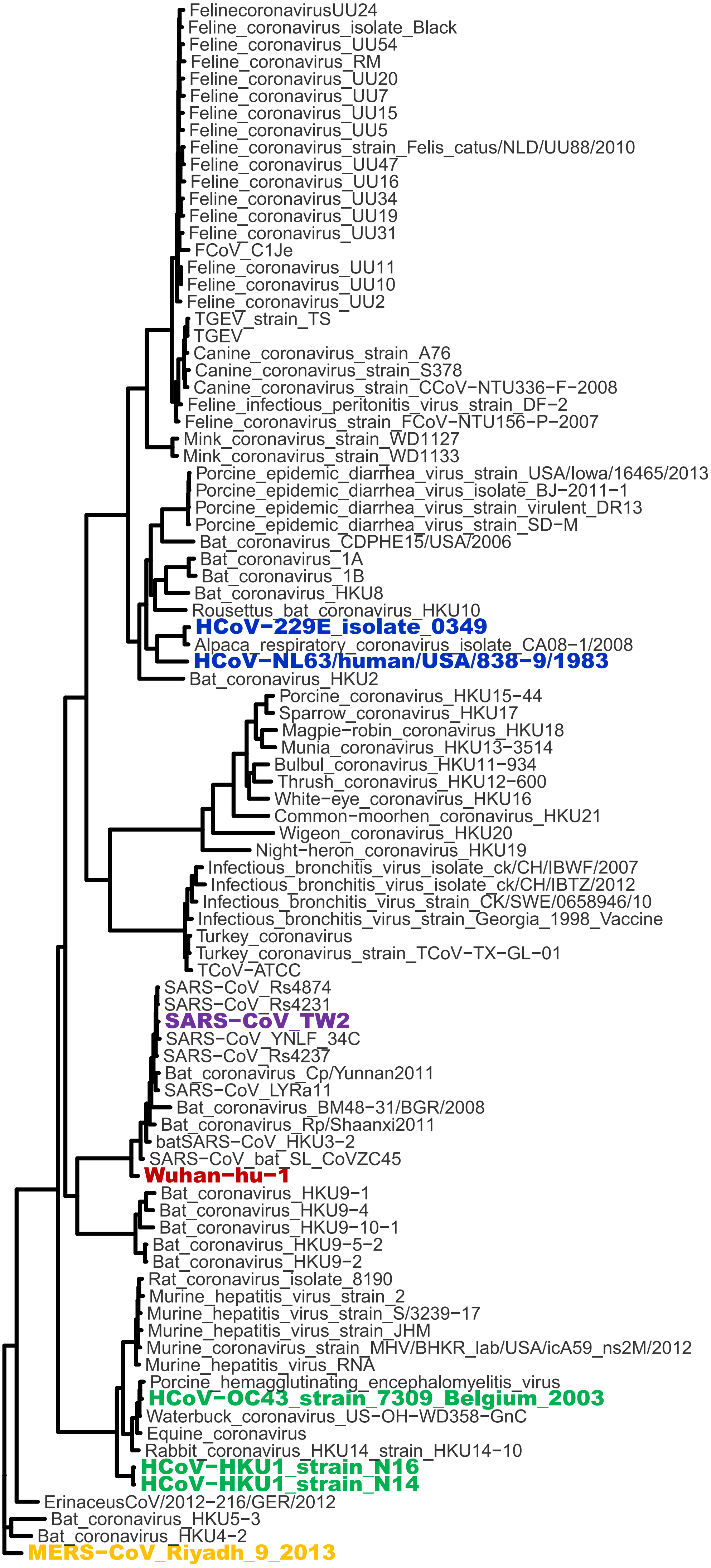

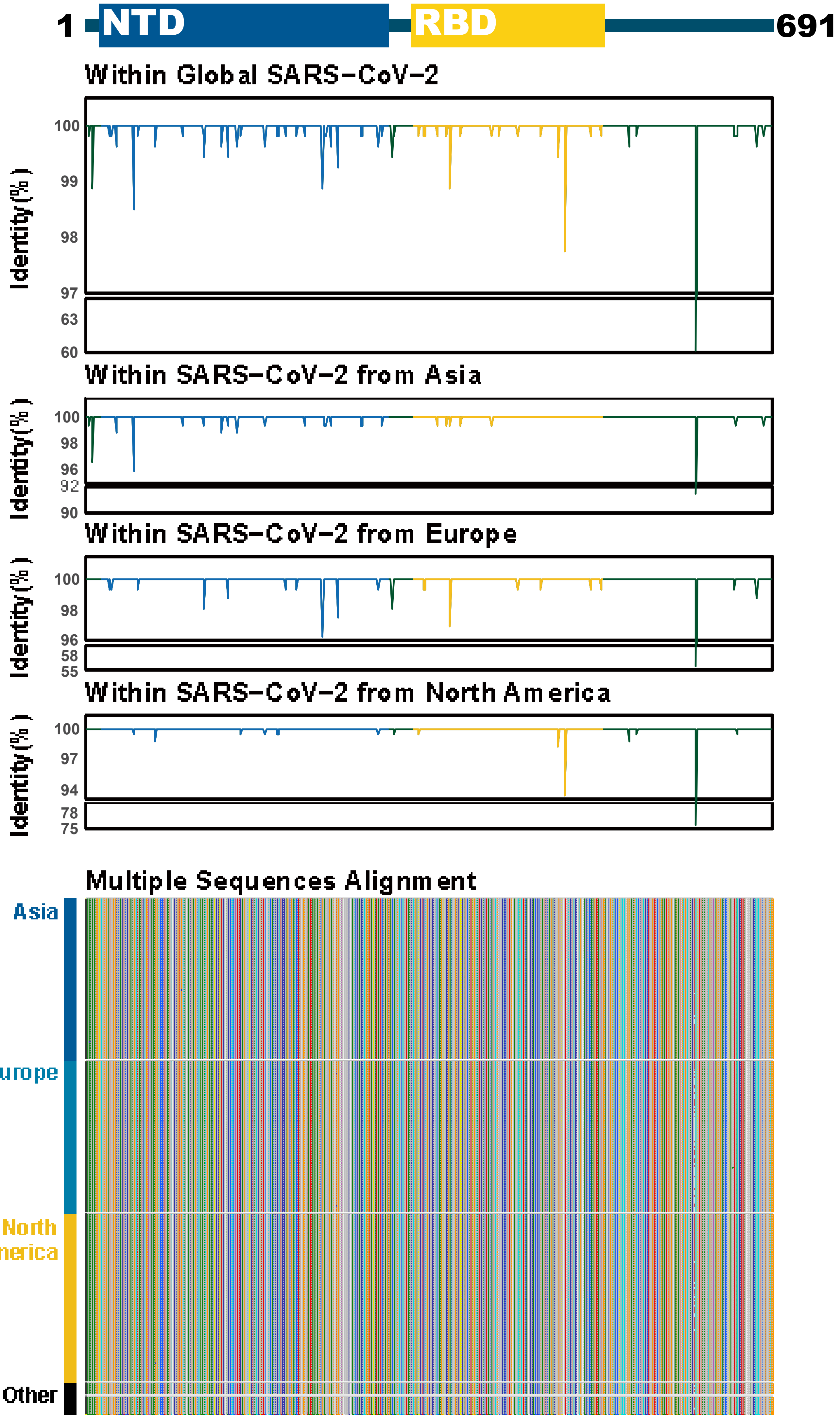

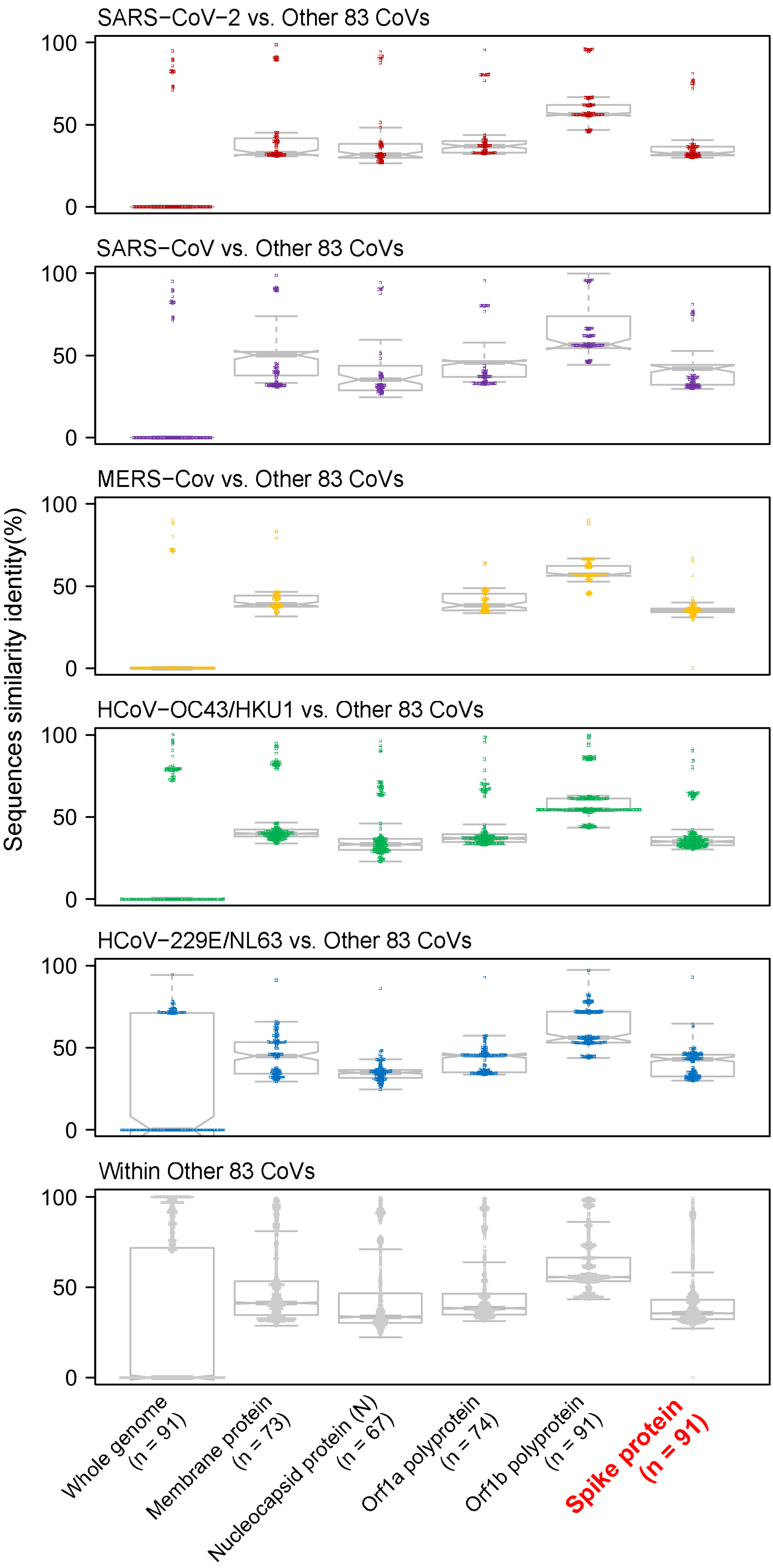

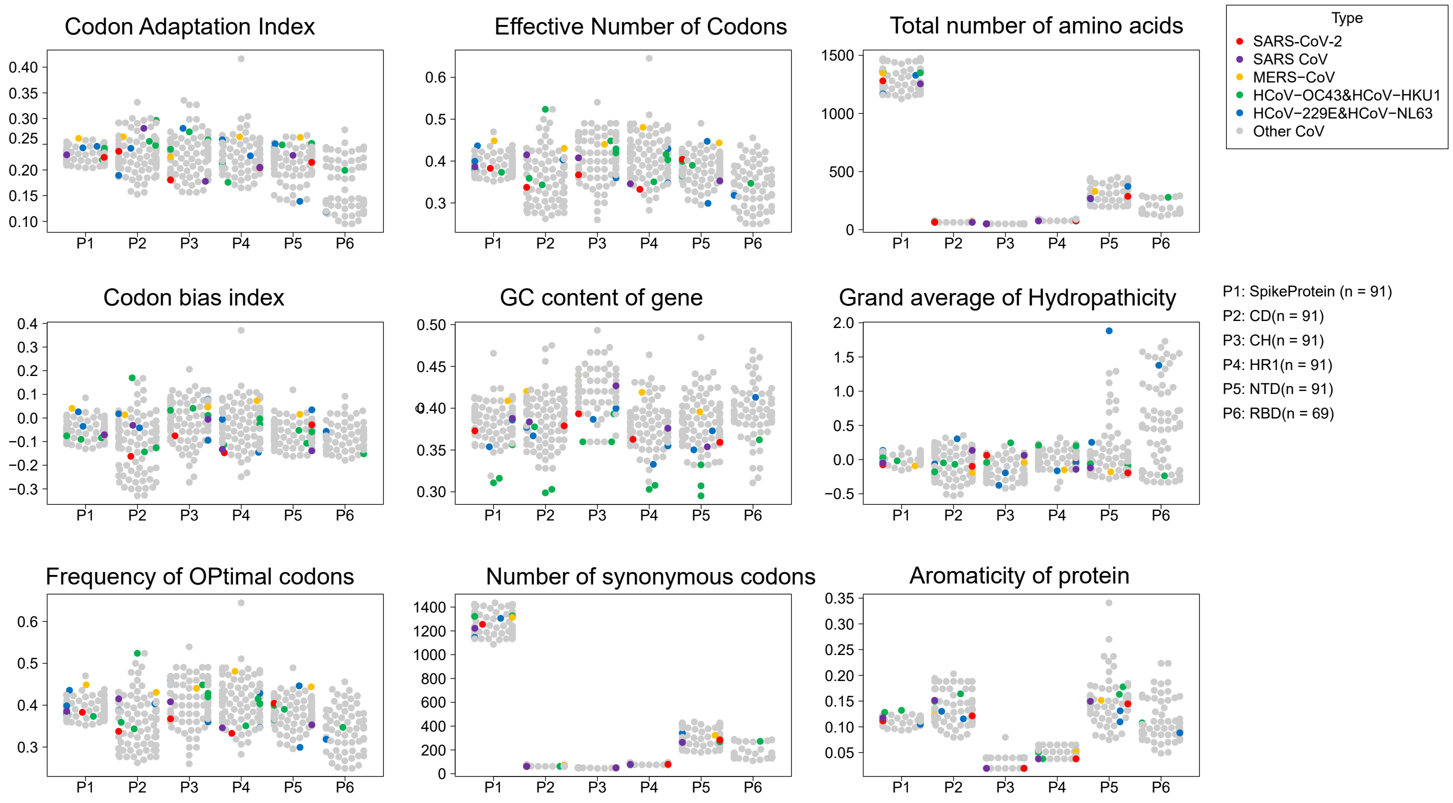

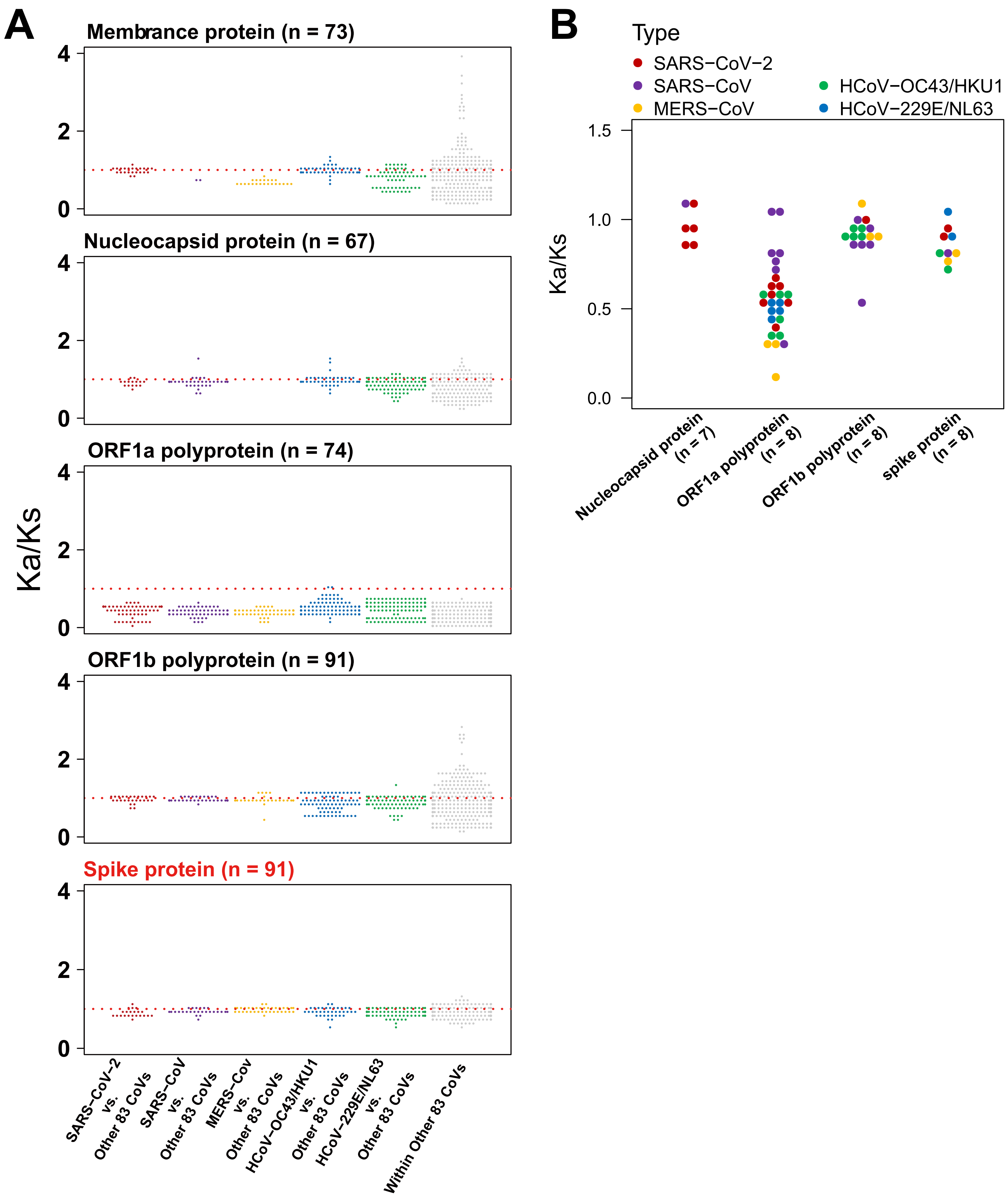

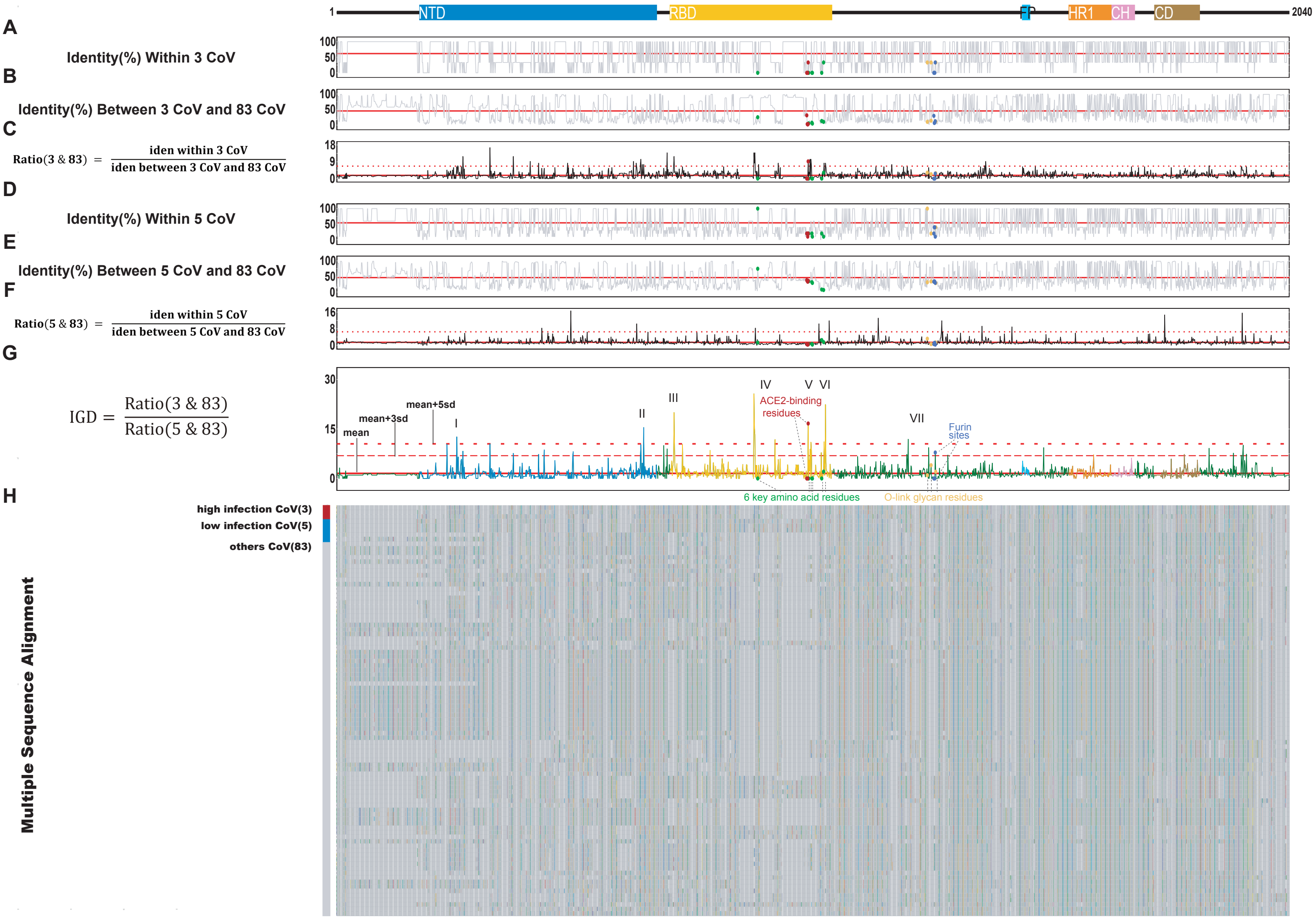
